# Neurofeedback helps to reveal a relationship between context reinstatement and memory retrieval

**DOI:** 10.1101/355727

**Authors:** Megan T. deBettencourt, Nicholas B. Turk-Browne, Kenneth A. Norman

**Author notes:** Corresponding author: Megan T. deBettencourt Institute for Mind and Biology 940 E 57^th^ St, Chicago, IL 60637.

## Abstract

Theories of mental context and memory posit that successful mental context reinstatement enables better retrieval of memories from the same context, at the expense of memories from other contexts. To test this hypothesis, we had participants study lists of words, interleaved with task-irrelevant images from one category (e.g., scenes). Following encoding, participants were cued to mentally reinstate the context associated with a particular list, by thinking about the images that had appeared between the words. We measured context reinstatement with fMRI, and related this to performance on a free recall test that followed immediately afterwards. To increase sensitivity, we used a closed-loop neurofeedback procedure, whereby higher levels of context reinstatement (measured neurally) elicited increased visibility of the images from the studied context onscreen. Our goal was to create a positive feedback loop that amplified small fluctuations in mental context reinstatement, making it easier to experimentally detect a relationship between context reinstatement and recall. As predicted, we found that higher levels of neural context reinstatement were associated with better recall of words from the reinstated context, and worse recall of words from a different context. In a second experiment, we assessed the role of neurofeedback in identifying this brain-behavior relationship by presenting context images again but manipulating whether their visibility depended on neural context reinstatement. When neurofeedback was removed, the relationship between context reinstatement and memory retrieval disappeared. Together, these findings demonstrate a clear effect of context reinstatement on memory recall and suggest that neurofeedback can be a useful tool for characterizing brain-behavior relationships.

**Abbreviated title:** Neurofeedback context

## Introduction

A key aspect of modern theories of context and memory (e.g., Polyn et al., 2009) is the ability to deliberately enact mental time travel: reinstating contextual features associated with a prior event in order to gain access to memories from that event (DuBrow et al., 2017; Manning et al., 2014). These theories predict that, following successful mental context reinstatement, memory performance should be improved for items encoded in the reinstated context relative to items from other contexts.

In this study, we set out to obtain neural evidence that deliberate context reinstatement predicts subsequent memory: We instructed participants to mentally time travel to a particular event and measured (based on brain activity) how well they did this, with the goal of showing that successful mental context reinstatement predicts successful recall of items from that context. Other related work has examined category-specific activation during a free recall task and found neural evidence for the to-be-recalled category prior to item recall (Polyn et al., 2005). However, this work confounded item and context activation, making it unclear whether the evidence for a category (e.g., face) reflected retrieval of a specific face item or a general face context.

To measure context recall separately from item recall, we used a method previously developed in our lab (Gershman et al., 2013), in which pictures of faces and scenes were used to establish contexts; both of these categories are known to robustly activate regions of visual cortex in fMRI (O’Craven et al., 1999). Specifically, we presented task-irrelevant pictures from one of these categories (either faces or scenes) interposed between to-be-learned word stimuli, thereby using these images to create a “context” for these words. Having established this item-context link, we were able to use neural activation of the “context” category to track whether participants were mentally reinstating the context. We have previously used this approach to predict memory misattribution errors (Gershman et al., 2013) and to explore intentional forgetting (Manning et al., 2016).

Despite the utility of this approach, context reinstatement is a subtle and dynamic internal mental process that is difficult to measure precisely. To amplify sensitivity to small neural fluctuations indicative of context reinstatement, we used a fMRI neurofeedback design (deBettencourt et al., 2015; Stoeckel et al., 2014; Sulzer et al., 2013). During time periods when participants were instructed to think back to a particular context (e.g., the list studied with interspersed scenes), we monitored in real time for neural activity relating to the context (e.g., scenes), while at the same time showing a stream of images from the target context (scenes that were actually presented during learning of the target word list) superimposed on images from the non-target context (faces that were presented with the other word list). When we detected brain activity relating to the target context, we increased the relative visibility of images from the target context. Participants were aware that the scene/face mixture proportion indicated their success at the context reinstatement task.

Our goal was to create a positive feedback loop where increased internal mental context reinstatement led to increased visibility of picture cues from the target context that triggered even more context reinstatement, thereby amplifying mental context reinstatement and (through this) boosting our ability to relate these neural fluctuations to memory performance. In our prior work (deBettencourt et al., 2015) we used a similar kind of neurofeedback to externalize participants’ top-down attentional state (i.e., whether they were attending to faces or scenes). Specifically, we instructed participants to attend to faces or scenes while they viewed superimposed faces and scenes; when their attention to the target category lapsed (as indexed by reduced category-specific evidence) we made that category less visible. The goal in that study was cognitive training, i.e., improving participants’ ability to detect and hence prevent attentional lapses. In the present study, the goal of neurofeedback was to amplify fluctuations in context reinstatement, not for the purpose of training participants, but rather to improve our ability as experimenters to detect these fluctuations and relate them to behavior.

We developed a task composed of three phases: encoding, context reinstatement, and recall (**Figure 1**). First, participants studied two lists of sixteen words; the first list (List A) was encoded in one of the category contexts (e.g., with scenes interleaved between the words) and the second list (List B) was encoded in the other category context (e.g., with faces interleaved between the words). After encoding (and a brief period of math distraction), participants were cued by the list name (e.g., List B) to reactivate the context associated with either the first or second list. Then, participants were presented with composite face/scene images, initialized to equal proportions (0.5/0.5) of each category. During context reinstatement, participants were instructed to think about the target category (e.g. faces), and were given real-time neurofeedback using the method described above. After context reinstatement, participants were presented with another list name (usually the list that had been cued, e.g., List B), which served as a memory probe. Their instructions were to freely recall as many words as possible from the probed list in any order. Participants’ vocal responses were recorded in the scanner during the recall phase.

**Figure 1.**
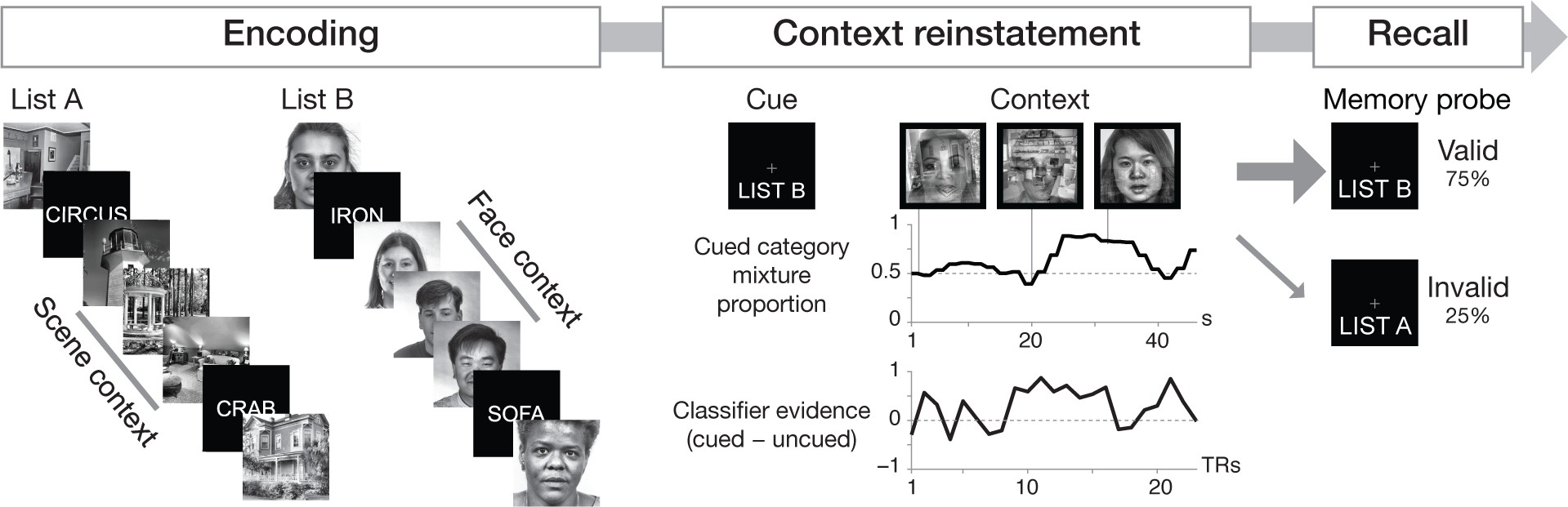
Study procedure. An example run of the task in Experiment 1. Each run was composed of encoding, context reinstatement, and recall phases. During the encoding phase, participants studied two lists of sequentially presented words, Lists A & B. Each of the lists was embedded in a different context by interleaving the words with images of a single category (scenes or faces). During the context reinstatement phase, participants were provided with a list name (e.g., List B) as a cue for which context (either scenes or faces) to reinstate. Participants were presented with composite face/scene images, initialized at 50% face and 50% scene. This mixture proportion was adjusted during the context reinstatement period to reflect the real-time decoding evidence for the cued context. The top row shows representative composite images. The middle row shows the corresponding proportion of the cued category of the composite image. The bottom row shows the real-time category evidence for the cued minus the uncued category for each TR during the context reinstatement period. Greater evidence for the cued context resulted in more of that category in the composite image (and less evidence resulted in less of that category). During the recall phase, participants were presented with a list name as a memory probe. Then, they were instructed to freely recall as many words as possible from the probed list. In validly cued runs (6 of 8 runs, 75%), the memory probe was the same as the context cue. In invalidly cued runs (2 of 8 runs, 25%), the memory probe was different from the context cue

Critically, in Experiment 1, we manipulated whether the context that participants were asked to reinstate matched the list they were subsequently asked to recall (e.g., reinstate the List B context, then recall List B; the *validly cued* condition) or whether the reinstated context mismatched the list they were asked to recall (e.g., reinstate the List B context, then recall List A; the *invalidly* cued condition). This manipulation was inspired by many studies of visual attention (e.g., Posner, 1980), which find that valid spatial cues improve performance and invalid cues impair performance. These findings are often explained in terms of spatial attention being focused on the cued location, which improves subsequent processing when the target appears at that location and impairs processing when the target appears elsewhere and attention needs to be reoriented. We expected an analogous effect in the memory domain with our context reinstatement manipulation, whereby the cue orients reinstatement towards the targeted list, improving recall from that list and impairing recall from other lists (Polyn et al., 2009). In order to ensure the effectiveness of the cueing procedure, cues were valid 75% of the time (6 of 8 runs). Invalidly cued runs (25%) occurred when the cue (e.g., List B) did not match the probe (e.g., List A). Our key prediction was that the relationship between target-category neural activity and recall behavior would be *positive* in the valid condition (i.e., greater reinstatement of the target context should help participants remember items from the target list) and *negative* in the invalid condition (i.e., greater reinstatement of the instructed context should be detrimental because participants were instructed to “mentally time travel” to the wrong context).

## Experiment 1: Materials and Methods

### Participants

Twenty-four adults participated in Experiment 1 for monetary compensation (14 female, 22 right-handed, mean age = 20.9 years). Power analyses could not be performed because of the use of a new paradigm and unknown behavioral and neural effect sizes. The sample size was chosen before the start of the experiment to match previous studies using a similar paradigm (Manning et al., 2016). Six additional fMRI participants were excluded from Experiment 1: two because of technical problems with real-time data or audio acquisition, one for falling asleep during several runs, and three for excessive motion, defined both within (≥ 3 mm) as well as across run (≥ 5 mm), due to the lack of real-time motion correction across runs during the fMRI session. All participants had normal or corrected-to-normal visual acuity and provided informed consent to a protocol approved by the Princeton University Institutional Review Board.

### Stimuli

*Word lists.* Prior to the experiment, we created 16 lists of words, with each list containing 16 words. Words and lists were derived from those used in a previous experiment (Manning et al., 2016). In Experiment 1, participants were presented with 16 lists in total. Each of these lists was randomly paired with a context of faces or scenes. The order of the words within lists and the order of lists during the experiment were randomized for each participant.

*Images.* Images consisted of grayscale photographs of male and female faces (Phillips et al., 1998) and indoor and outdoor scenes (Xiao et al., 2010). These images were combined into composite stimuli by averaging pixel intensities using various weightings (e.g., 60% face, 40% scene). The stimuli were displayed on a projection screen at the back of the scanner bore and viewed with a mirror attached to the head coil.

### Procedure

*Localizer runs.* Participants completed two localizer runs, viewing blocks of scene, face, and object images. Each block consisted of 12 images, presented for 1s with a 0.5s period of fixation between each image. A total of 12 blocks were presented, with 4s of fixation separating each block. Participants were asked to detect back-to-back image repetitions, and respond by pressing a button.

*Memory runs.* Each memory run began with 38s fixation, followed by three phases: study, context reinstatement, and recall. During the study phase, participants studied two lists. For each list, the name of the list (either “LIST A” or “LIST B”) was presented for 3s, followed by 1.5s fixation. Each word was presented for 3s. Between each word, 3 images (either faces or scenes) were presented, each for 1s. In total, each list was composed of 16 words and 45 images, with 6s of fixation at the end. For each memory run, one studied list was paired with a scene context and the other with a face context. After the study phase, there was a brief period of math problems (15s) to distract participants and prevent rehearsal: the response mapping for the math problems was first displayed for 1.5s, followed by 9 math problems for 1.5s each. Each math problem involved determining whether the sum of two single digit numbers (e.g., 3+4) was even or odd. After the math section was complete, there was a 3s fixation period.

At the start of the context reinstatement period, the cue (either “LIST A” or “LIST B”) was displayed for 3s. This was followed by 33.75s of 45 images, each displayed for 0.75s. For feedback runs, these images were composite face and scene images. The first two images (1.5s) were always initialized at 50% face and 50% scene. The mixture proportions for the remaining trials were determined on the basis of real-time multivariate pattern analysis (MVPA) of the fMRI data, ranging from 17% to 98% of the category of the cued context (83% to 2% of the category of the uncued context) as in (deBettencourt et al., 2015). In Experiment 1, the images presented during context reinstatement were composites of the actual faces and scene images that had appeared during the study phase for the current memory run. Both the face and scene images appeared in a random order. At the end of the context reinstatement period, before the memory probe appeared, there was a 0.75s fixation period.

The recall phase began with a 3s memory probe, indicating the name of the list to recall (either “LIST A” or “LIST B”). Then, the fixation dot on the screen turned green to indicating the start of the recall period. Participants were given 45s to recall the items from one of the lists in any order. At the end of the recall period, the fixation dot turned back to white for 4s. Then, participants were presented with a point score from that run’s context reinstatement period, corresponding to classifier decoding performance during the context period (i.e., classifier accuracy for the target category). No feedback was provided on their recall performance. At the end of the experiment, participants received up to $10 extra corresponding to their cumulative scores on all the runs.

Participants completed 8 runs of the task (6 valid and 2 invalid). There were more valid than invalid runs, to ensure that participants would be motivated to attend to the cue. In addition, the first two runs of the experiment were always valid. One invalid run occurred during runs 3–5, and the other invalid run occurred during runs 6–8, and the invalid runs were not permitted to occur back-to-back (i.e., runs 5 and 6). Behavioral and neural analyses relating to recall were conducted on runs where it was possible an invalid cue could occur (i.e., runs 3–8). Runs 1 and 2 were excluded so as to minimize the temporal imbalance and any resulting practice effects between valid vs. invalid conditions. The two invalid runs were counterbalanced for list cue (one invalid run cued List A and the other cued List B) and cued context category (one invalid run cued the scene category and the other cued the face category).

Importantly, participants were aware that what they saw onscreen during the reinstatement period was controlled by their brain activity. Before the fMRI session, they were given instructions about the experiment, which included the feedback manipulation. They were told that the images in the context reinstatement period would reflect our measurements of their mental context, and that the images would get easier to see if they were reinstating context well. They completed an abbreviated run of the task outside the scanner to familiarize themselves with the experimental design. During that run, they were shown examples of a composite stimulus, and how the mixture proportion could change due to our measurements.

### Data acquisition

Experiments were run using the Psychophysics Toolbox for Matlab (Brainard, 1997; Pelli, 1997). Neuroimaging data were acquired with a 3T MRI scanner (Siemens Skyra) using a 20-channel head and neck coil. We first collected a scout anatomical scan to align axial functional slices to the anterior commissure-posterior commissure line. Then, a high-resolution magnetization-prepared rapid acquisition gradient-echo (MPRAGE) anatomical scan was acquired to use for real-time spatial registration. Functional images were acquired using a gradient-echo, echo-planar imaging sequence (1.5s repetition time or TR, 29 ms echo time, 3 × 3 × 3.5 mm voxel size, 64 × 64 matrix, 192 mm field of view, 27 slices).

### Experimental design and statistical analysis

Because some of the data violated the assumption of normality, all statistics were computed using a nonparametric random-effects approach in which participants were resampled with replacement 100,000 times (Efron and Tibshirani, 1986). Null hypothesis testing was performed by calculating the proportion of the iterations in which the bootstrapped mean was in the opposite direction. One-sided tests were used for directional hypotheses and two-sided tests for non-directional hypotheses. Correlations between two variables were estimated with Spearman’s rank correlation after applying robust methods to eliminate the disproportionate influence of outliers in small samples (Pernet et al., 2013). Outliers were excluded only if they exceeded 2.5 standard deviations from the mean; all outlier exclusions are noted in the text.

### Real-time analyses

*Preprocessing.* At the start of the fMRI session, an anatomical region-of-interest (ROI) was registered to the native functional space using FSL’s FLIRT. For Experiment 1, the temporal lobe was *a priori* selected to be the ROI given its known involvement in representing contexts for memories. During the fMRI session, functional data were reconstructed and prospective acquisition correction and retrospective motion correction were applied. After motion correction, the file was transferred to a separate analysis computer for the remainder of the real-time analyses. The anatomical ROI mask was applied to reduce the voxel dimensionality. The volume was spatially smoothed in Matlab using a Gaussian kernel with full-width half-maximum (FWHM) = 5 mm. In Experiment 1, a high-pass filter adapted from FSL (cutoff = 100s) was applied in real time. After each localizer run, the BOLD activity of every voxel was z-scored over time. During memory runs, the BOLD activity of each voxel was *z*-scored starting after the study period based on the mean and standard deviation until then.

*Multivariate pattern analysis.* A multivariate pattern classifier was trained on data from the face and scene blocks from both localizer runs. Labels were shifted 3 TRs (i.e., 4.5s) forward in time to account for the hemodynamic lag. For Experiment 1, we conducted MVPA using penalized logistic regression with L2-norm regularization (penalty = 1).

The trained model was tested in real time on brain volumes obtained during the context reinstatement period. For each volume, the classifier estimated the extent to which the brain activity pattern matched the pattern for the two categories (face and scene) on which it was trained (from 0 to 1); we will refer to this quantity as *classifier evidence*. The neurofeedback was based on the difference of classifier evidence for the task-relevant category minus task-irrelevant category. However, given that this was a binary classifier, the evidence for each category is anticorrelated. The subtraction means that the difference in evidence ranges from −1 (complete evidence for the wrong category) to 1 (complete evidence for the correct category).

*Neurofeedback*. The output of the classifier was used to determine the proportion of the images from the cued and uncued categories in the composite stimulus on the next trial. The preprocessing and decoding of volume *i* were performed during volume *i*+1 and the classifier output was used to update the stimulus mixture for the two trials in volume *i*+2. This resulted in a minimum lag of 1.5s (one TR or two composite images) between data acquisition and feedback. Moreover, classifier output was averaged over a moving window of the preceding three volumes (*i*−2, *i*−1 and *i* for feedback in volume *i*+2), meaning that feedback was based on brain states 1.5–6s in the past. The average classifier output was mapped to a proportion of the task-relevant category using a sigmoidal transfer function (deBettencourt et al., 2015).

### Behavioral analyses

Vocal responses were recorded during the recall period using a customized MR-compatible recording system (FOMRI II; Optoacoustics Ltd.). We used the Penn TotalRecall tool to score and annotate the audio. All annotations were completed without knowledge of the experimental design and were verified by an independent scorer who had no knowledge of the experimental manipulation or hypotheses.

### Decoding accuracy

Multivariate pattern classifiers were trained using the face and scene blocks from both localizer runs. Classifier performance was assessed by testing the classifier on TRs during the context reinstatement period (as with classifier training, labels were shifted forward 3 TRs (4.5s) to account for hemodynamic lag). Two measures were used: classifier evidence for the cued category and accuracy. A TR was labeled as accurate if the evidence for the cued category was greater than the evidence for the uncued category. To assess whether there was any bias at the start of the context reinstatement period, the classifier accuracy was calculated for the data from the last TR during cue presentation (i.e., when participants were being informed which list to reinstate). To evaluate classifier performance during the context reinstatement period, accuracy was computed during the entire context reinstatement phase. Chance was assessed by permuting labels 100,000 times and recalculating classifier accuracy for each of these permutations.

### Relationship to behavior

To explain how context reinstatement related to memory behavior, we first obtained summary measures for each of these components for each run: (1) how successful participants were at reinstating the cued context and (2) their memory performance. Context reinstatement was operationalized as the classifier evidence for the cued context (relative to the uncued context) during the period of context reinstatement. To ensure that we were maximally sensitive to the period of time that context evidence was measurable, we first identified the time points where on average, across participants, there was significantly more evidence for the cued (vs. uncued) category, and we limited our per-run measurement of context reinstatement to these TRs. Memory performance was calculated as total number of distinct words that were recalled for the probed list.

## Experiment 1 Results

### Real-time multivariate decoding of mnemonic context

We first assessed the overall degree to which participants were reinstating the cued context during the reinstatement period; we expected that there would be greater classifier evidence for the cued context, compared to the uncued context. Consistent with this prediction, the real-time multivariate pattern classifier reliably decoded the cued context during the context reinstatement period (mean accuracy = 0.58, 95% CIs 0.57–0.60; chance = 0.5; *p* < 0.00001; **Figure 2**). At the start of the context reinstatement period, there was no reliable evidence for the cued category (mean evidence = 0.52, 95% CIs 0.43–0.58; chance = 0.5; *p* = 0.63, two-tailed).

**Figure 2.**
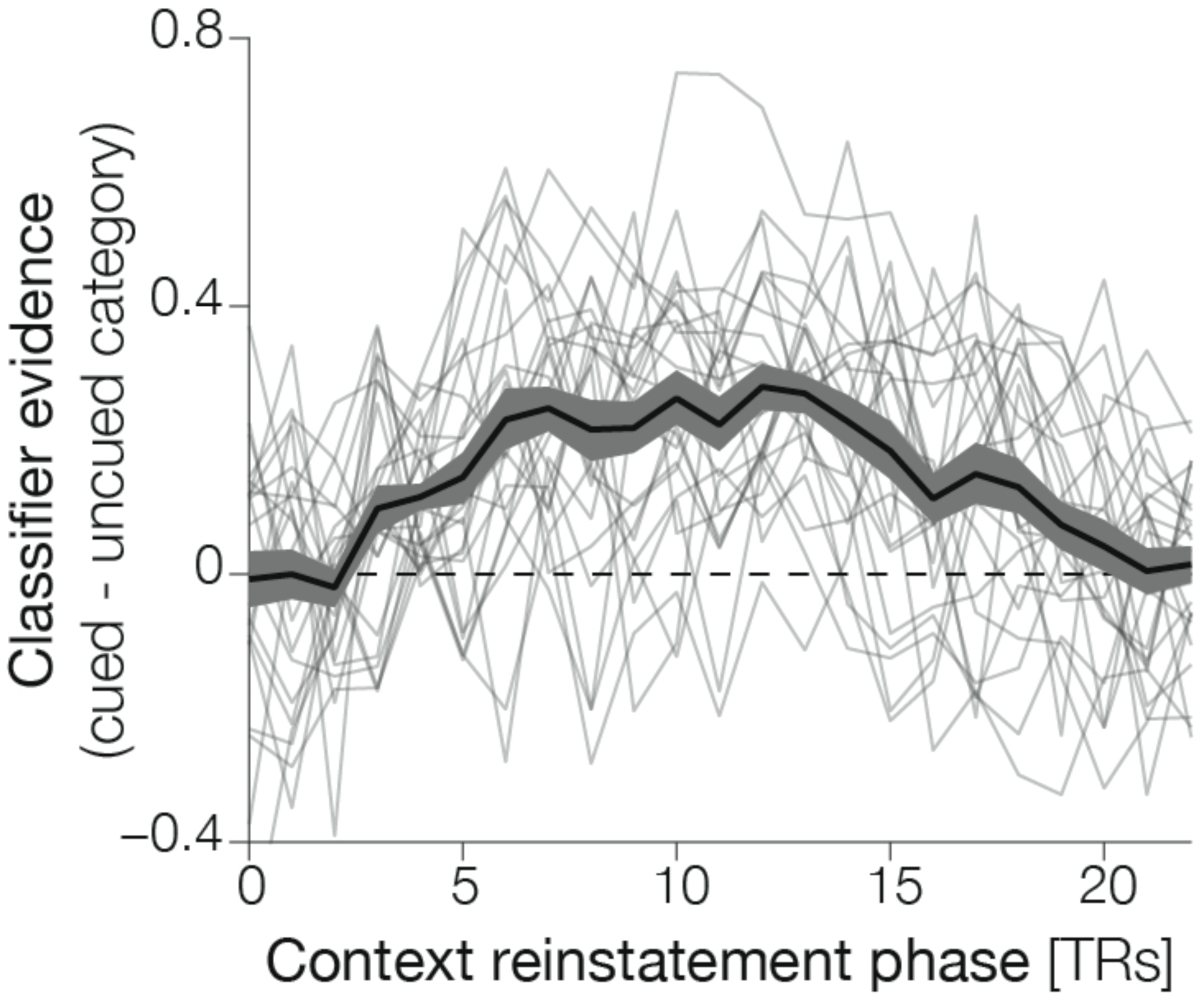
Timecourse of real-time multivariate classifier decoding of context. The average classifier evidence for each participant across all feedback runs is plotted in thin gray lines. The average timecourse across participants is plotted in black, with the gray ribbon indicating the standard error of the mean. The *y-axis* shows the classifier evidence for the cued category minus the classifier evidence of the uncued category. The *x-axis* shows the number of TRs (1.5s) during the context reinstatement phase.

### Behavioral effects

During the validly cued runs of Experiment 1, participants were cued to mentally reinstate the same context for the list they were later asked to recall. During the invalidly cued runs of Experiment 1, the context reinstatement cue did not match the memory probe. If prior context reinstatement influenced later memory, overall memory recall should be higher on valid versus invalid runs. Consistent with the idea that reinstating an appropriate context boosts memory recall, more items were recalled in the valid as compared to the invalid condition (M*_valid_* = 5.85, 95% CIs 4.83–7.14; M*_invalid_* = 5.29, 95% CIs 4.08–6.75; *p* = 0.031; **Figure 3a-c**).

**Figure 3.**
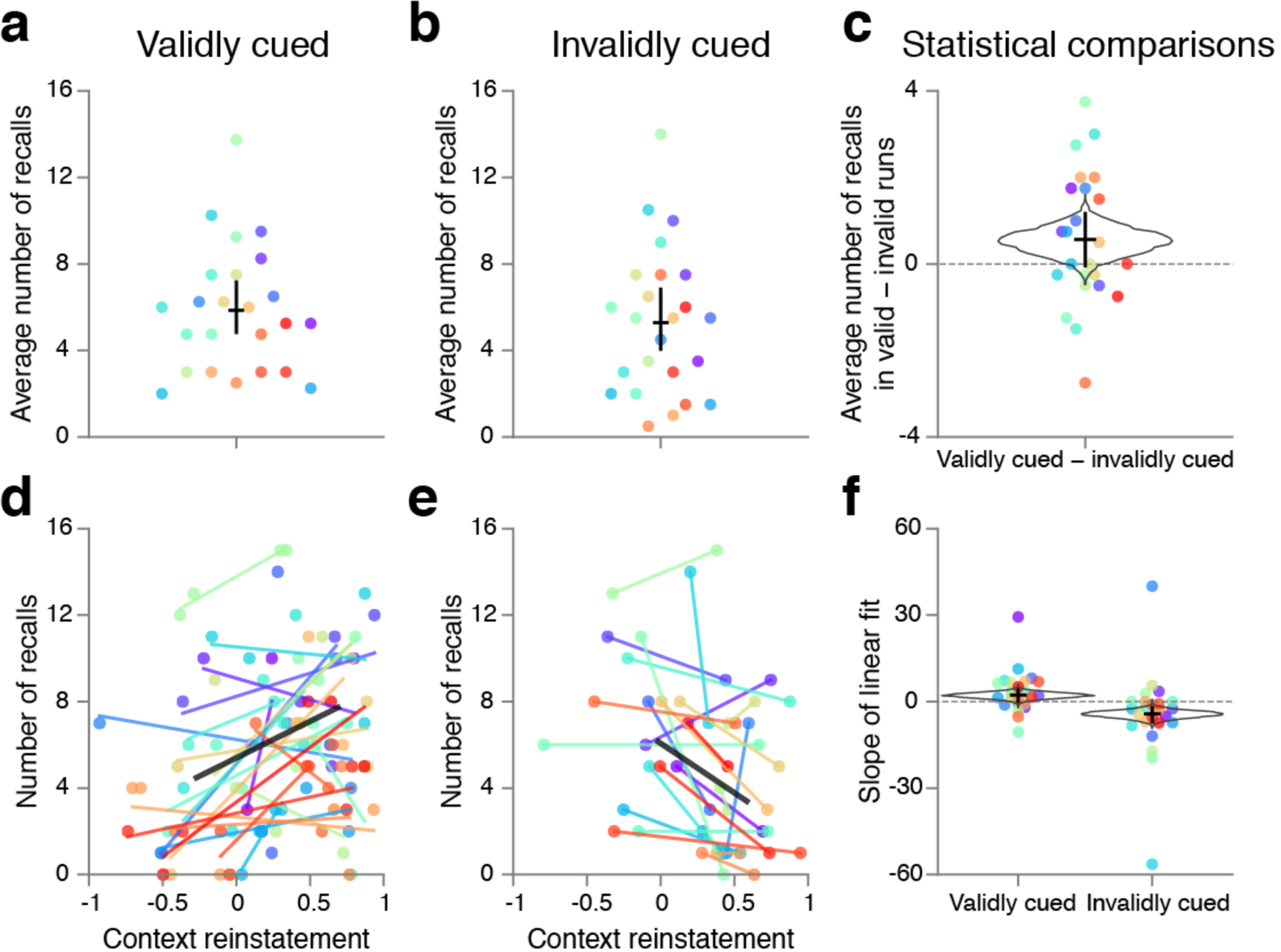
Effects of context reinstatement cue validity on memory in Experiment 1. (a) Memory recall performance for validly cued feedback runs, where the cue at the start of the context reinstatement period matched the memory probe at the start of the free recall period. Each colored dot indicates the average number of words recalled per participant (n=24), and individual participants are depicted in unique colors. The horizontal black line indicates the population average, and the vertical black line indicates 95% confidence intervals. (b) Memory recall performance for invalidly cued feedback runs. (c) Memory recall performance difference based on condition (valid minus invalid). The grey outline is the resampled histogram for the nonparametric statistical tests. Memory performance was enhanced following valid cues (*p* < 0.05). (d) In validly cued runs, the amount of context reinstatement positively predicted the number of recalls. There was a reliably positive relationship (*p* < 0.05) between the context reinstatement, that is the evidence for the cued context minus uncued context (*x-axis*) and the total number of recalls (*y-axis*) for each run. The linear fit across runs within a single participant is depicted in each participant’s unique color. The mean linear fit is depicted in black. (d) In invalidly cued runs, the amount of context reinstatement negatively predicted the number of recalls. There was a reliably negative relationship (*p* < 0.00005) between the context reinstatement (*x-axis*) and the total number of recalls (*y-axis*) for each run. (e) Statistics were computed using the slopes of the linear fits per condition. Each dot corresponds to the slope of the linear fit in a single condition (valid or invalid) for each participant.

The key question that we asked in this study pertains to the *relationship* between neural context reinstatement and memory behavior: We hypothesized that higher levels of neural context reinstatement would correlate with higher levels of recall for validly cued lists and lower levels of recall for invalidly cued lists.

To test this hypothesis, we computed the relationship between neural context reinstatement and recall performance across runs (within participants, separately for valid and invalid runs) and then averaged this measure across participants. Each participant had 6 valid memory runs; for each run, we computed our index of context reinstatement (classifier evidence for the cued vs. uncued context over the course of the reinstatement period), and also the total number of recalls for each tested list. This yields a participant-specific scatterplot with 6 points (one per memory run). For each participant, we computed the slope of the line relating context reinstatement to recall performance. We then evaluated the reliability of the slope of this line across participants. For invalid memory runs, we used the same analysis procedure (this time focusing on the 2 invalid runs) to estimate the relationship between context reinstatement and recall performance on these runs.

As predicted, we observed a reliably positive relationship between neural context reinstatement and recall performance across valid runs (slope*_valid_* = 2.25, 95% CIs 0.09–4.07; *n* = 24; *p_valid_* = 0.015; **Figure 3d&f**). Thus, greater amounts of context reinstatement resulted in better memory. For invalid runs, we expected there to be a reliably negative relationship between neural context reinstatement and recall performance, and this was also upheld (slope*_invalid_* = −4.24, 95% CIs −7.14– −2.02; *p_invalid_* = 0.00009; n = 22, 2 outliers excluded; **Figure 3e&f**). Importantly, the relationship between context reinstatement and memory was significantly most positive for valid than invalid runs (*p_diff_* = 0.01; **Figure 3f**). As noted earlier, these correlations were completed using data from the TRs where classification was reliably above chance (in the group analysis) for maximal power within each category. However, the observed effects were insensitive to the exact selection of time points: The same results were observed when using data from the entire context reinstatement period (slope*_valid_* = 2.75, 0.85–5.59, *p_valid_* = 0.004; slope_invalid_ = −3.43, −6.95–0.17; *p*_invalid_ = 0.005; slope*_diff_* = 7.16, 2.24–11.91, *p*_diff_ = 0.009).

Note that the valid-condition results, considered on their own, could be explained in terms of a third factor (e.g., general alertness) that positively affects both context reinstatement and recall performance. However, the valid and invalid results can not *together* be explained this way: If general alertness benefits both context reinstatement and recall, resulting in a positive relationship between them, this relationship should be observed in *both* the valid and invalid conditions, but this was not the case. The most parsimonious account of the valid and invalid results together is our hypothesis, that context reinstatement facilitates recall of a one list at the expense of the other.

For the analyses reported above, context reinstatement and memory were calculated across the valid and invalid runs separately within participants. However, the validity of the cue was fairly high (75%) and therefore the number of invalid runs was low (2 invalid runs in total, during runs 3–8). When participants did not differ substantially in context reinstatement for these two runs, this resulted in extreme slope values. We wanted to be certain that the relationship between context and memory in the invalid runs was not driven by any extreme values and/or our outlier exclusion procedure. Therefore, we conducted a similar analysis, but computing the relationship at the group, rather than individual, level. First, to keep the analysis focused on within-participant variance (as opposed to between-participant variance), we normalized neural context reinstatement scores and number of recalls within condition (valid, invalid) for each participant. Then, we calculated the linear relationship (i.e., slope) between context reinstatement and memory recall performance across all invalid runs from all individuals. The relationship between context reinstatement and memory recall performance remained negative (slope*_invalid_* = −0.54, −0.83– −0.21). We assessed the reliability of this relationship by conducting a bootstrap correlation analysis in which we resampled participants with replacement and calculated the correlation on each new sample (Kim et al., 2014). This bootstrapped correlation was reliably negative (*p_invalid_* = 0.001). These results provide additional evidence that reinstating an inappropriate context with neurofeedback prior to recall adversely influences subsequent memory performance.

These same group-wise analyses were conducted by normalizing context reinstatement and recalls within the validly cued runs. We found that the slope computed across valid runs from all participants remained positive (slope*_valid_* = 0.35, 0.10–0.54). A bootstrapped correlation was calculated by resampling participants with replacement and calculating the correlation on each new sample; this bootstrapped correlation was reliably positive (*p_valid_* = 0.002). Lastly, with these group-wise analyses the difference between the bootstrapped correlations in the valid and invalid conditions remained robustly different (*p_diff_* = 0.00012).

### Simulations

To summarize the results thus far: In Experiment 1, we obtained a relationship between mental context reinstatement (measured neurally) and subsequent recall, using neurofeedback. This raises the question: How important was the use of neurofeedback in obtaining these results? Could we have obtained this relationship between mental context reinstatement and subsequent recall without using neurofeedback? As discussed earlier, we used neurofeedback in Experiment 1 because of our intuition that it improves our ability to measure subtle fluctuations in context reinstatement, compared to other approaches. To verify this intuition, we ran simulations comparing our neurofeedback condition to various other (non-neurofeedback) control conditions. The goal of these simulations was to consider potentially informative control experiments that could be run.

The simulations sought to capture what occurs during the context reinstatement period of our experiment (**Figure 4a**). During the first 2 (simulated) TRs, a top-down cue biased the mental context towards a particular category (scene). Internal context was first established by that cue, plus additional noise, with the strength of internal context reinstatement varying across runs of the simulation. Internal context, plus noise, determined classifier decoding. In the (simulated) neurofeedback condition, classifier decoding influenced perceptual evidence, which in turn, influenced internal context. At the end of the context reinstatement period, a random value (0–1) was assigned to each of the 16 words. The average level of internal context activation during the reinstatement period determined the threshold, and all words that exceeded this threshold were considered recalled. The correlation between decoding accuracy and number of words recalled was calculated.

**Figure 4.**
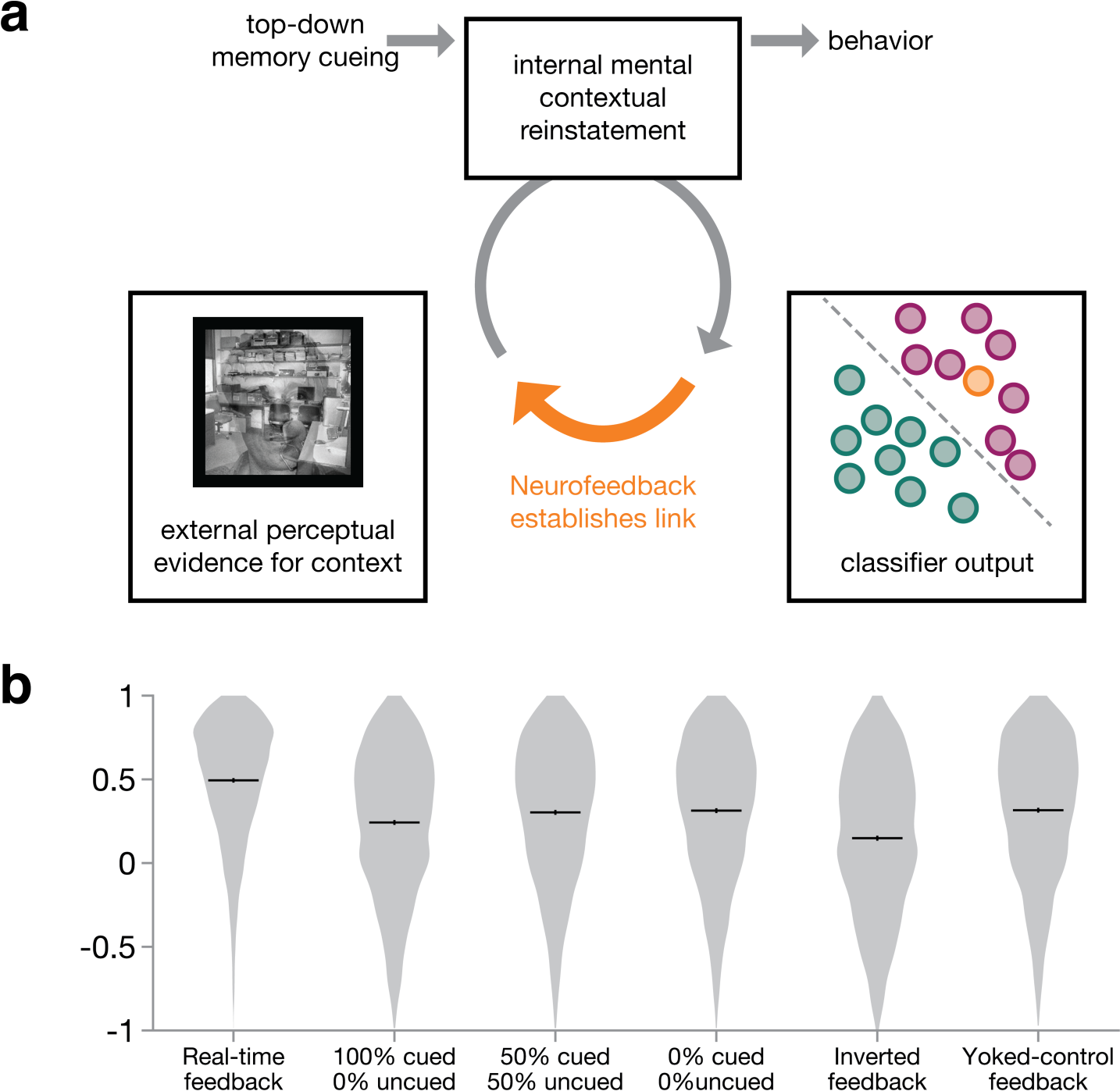
Simulating feedback. (a) A schematic of the hypothesized context reinstatement process with the link mediated by real-time neurofeedback in orange. (b) Results of computational simulations of the correlation between internal mental context and memory recall behavior. Simulations were completed for various manipulations of the feedback as well as perceptual evidence: real-time neurofeedback (in which the perceptual evidence reflects the classifier output of the cued category), maximal perceptual input (100% cued category, 0% uncued category), balanced perceptual input (50% cued category, 50% uncued category), no perceptual input (0% cued, 0% uncued), inverted real-time neurofeedback (in which the perceptual evidence reflects the classifier output of the uncued category), and yoked-control feedback (in which the perceptual evidence reflects the classifier output from another run. Each violin plot represents the correlations computed across 10,000 simulations. The mean correlation is depicted in the horizontal black line, and 95% CIs in the vertical black line.

We simulated five control conditions to contrast against real-time neurofeedback: First, we included controls where we held the amount of perceptual evidence for the cued category constant (at three different levels: 100%, 50%, 0%). In addition, we included two types of time-varying feedback: inverted feedback, in which the mapping between classifier evidence and perceptual evidence was flipped, and yoked-control feedback, in which the perceptual evidence was selected from a different run. By re-running these simulations 10,000 times, we established distributions of correlations between context and memory recall across these conditions.

## Simulation results and discussion

As shown in Figure 4b, the correlation between context reinstatement and recall observed in the feedback condition was higher than the other conditions (Spearman rank correlation: *r_feedback_* = 0.49, *r_100%cued_* = 0.24, *r_50%cued_* = 0.30, *r_0%cued_* = 0.31, *r_inverted_* = 0.15, *r_yoked_* = 0.32). Taken together, these results validate our intuition that closed-loop neurofeedback can positively amplify subtle fluctuations in the internal state of context, in comparison to many other possible control conditions.

In Experiment 2, we wanted to verify experimentally that the neurofeedback condition is especially sensitive to the relationship between context reinstatement and recall. In an ideal world, we could run all controls, but we only had the time and resources to focus on one. Choosing a control condition is not easy for neurofeedback studies, insofar as there are many different alternative hypotheses and each control condition addresses a subset of these hypotheses. For example, the yoked-control approach (which we used in deBettencourt et al., 2015) has several benefits: It controls for the specific stream of images that participants view, and it also controls for instructions provided to participants (both neurofeedback participants and yoked controls are told that their brain activity is controlling stimulus visibility, which should control for any general motivational effects of being told that you are in a neurofeedback experiment). However, it also has several drawbacks: If we find a worse relationship between brain activity and recall, it could be due to lack of accurate neurofeedback or because participants get distracted or frustrated when the feedback does not match their own sense of their mental state. That is, any observed differences between conditions might be due to yoking being harmful rather than neurofeedback being helpful. Also, if we yoke the images but do not say that fluctuations in visibility are due to neurofeedback, this fails to control for nonspecific motivational effects of participants thinking they are getting brain-based feedback (also, images varying in a way that has nothing to do with participants’ brain state might be distracting).

We next considered a control condition where pictures from the target context are 100% visible during the reinstatement period (instead of being mixed with pictures from the other controls). This control tests whether merely showing “reminder” images from the target context is sufficient to reveal a relationship between neural context reinstatement and recall behavior. If this is the case, then showing *fully visible* images from the target context should yield an especially robust effect. This control also has obvious drawbacks: It does not control for motivation that comes from participants thinking they are getting neurofeedback, and it does not match the exact image stream seen by neurofeedback participants. However, because there is no one perfect control, we decided to try this 100% visible control condition — showing images from the target context seemed to us to be the most direct way to trigger context reinstatement. In the discussion, we talk about inferences that we can (and cannot) glean from this condition.

## Experiment 2: Materials and Methods

Here we compare performance with feedback to a control condition in which stimulus information was not modulated by the participant’s mental context. We kept the basic structure of the runs the same as it was in Experiment 1, but eliminated the second list (participants only studied one list per run before recall) and removed the invalid condition in order to focus on the difference between valid neurofeedback and the control condition. For 6 of the 12 runs of Experiment 2, context reinstatement was provided as in the first experiment, and participants received neurofeedback with composite face/scene images as in Experiment 1. For the other 6 non-feedback runs, participants viewed all of the images that had appeared between words during the encoding phase at full coherence without any competitive information on the screen (100% cued context). That is, they viewed fully coherent scenes during the context reinstatement phase, rather than composite face/scene pairs, in order to provide the strongest possible visual cues for context reinstatement.

### Participants

Twenty-four adults participated in Experiment 2 for monetary compensation (11 female, 22 right-handed, mean age = 19.3 years). The sample size was matched to that of Experiment 1. Five additional fMRI participants were excluded from Experiment 2: one due to lack of understanding the instructions, and four for excessive motion, using the same standards as in Experiment 1. All participants had normal or corrected-to-normal visual acuity and provided informed consent to a protocol approved by the Princeton University Institutional Review Board.

### Stimuli

The word list stimuli for Experiment 2 were a subset of those used in Experiment 1: we selected 12 (of the original 16) word lists. The word lists were interleaved with scene images (not faces). During the context reinstatement portion of the feedback runs, face images (not presented at study) were overlaid on the studied scenes in order to alter the mixture proportions.

### Procedure

Participants completed two localizer runs, as in Experiment 1. Also as before, each memory run was composed of three phases: study, context reinstatement, and recall. The details of the memory runs were the same as in Experiment 1, except for the following changes: In Experiment 1, participants studied two lists in each memory run (one with faces and one with scenes); in Experiment 2, there was only a single list in each study period (List A), which was always paired with a scene context. Also, in Experiment 2, all lists were validly cued (i.e., the reinstated context always matched the list that participants were asked to recall). In Experiment 2, 6 of the runs (50%) were real-time neurofeedback runs and 6 of the runs (50%) were control, non-feedback runs. Participants were informed ahead of time that some runs would be feedback runs and some would not. These runs were counterbalanced such that every 4 runs (i.e., 1–4, 5–8, and 9–12) contained 2 valid feedback runs and 2 valid non-feedback runs. As such, there was no temporal imbalance, and all runs were included in the subsequent analyses. In the feedback runs of Experiment 2, the images presented during context reinstatement were composite face and scene images (as in feedback runs from Experiment 1). The scene images were those that had appeared during the encoding phase in a random order. The face images were randomly selected from a list without replacement so that each face image was only presented once throughout Experiment 2. In non-feedback runs during Experiment 2, the images presented during context reinstatement were scene images (fully coherent, or 100%). The scene images were those from the study phase, presented in random order.

### Data acquisition, statistical analysis, real-time analysis

Data acquisition and (offline) statistical analysis methods were identical to the methods used in Experiment 1. The real-time analysis methods were mostly the same as in Experiment 1, except for the following changes: First, in Experiment 2, the temporal lobe ROI that we used in Experiment 1 was expanded to also include the occipital lobe. This decision was based on offline analyses of the data from Experiment 1 that demonstrated that including the occipital lobe resulted in higher overall classifier accuracy. Second, in Experiment 1, a high-pass filter adapted from FSL (cutoff = 100s) was applied in real time, but in Experiment 2, no such high-pass filter was applied (since the run length was shorter). Third, in Experiment 1, we conducted MVPA using penalized logistic regression with L2-norm regularization (penalty = 1), but in Experiment 2 (based on the results of offline reanalysis of data from Experiment 1 to optimize classification), we modified the logistic regression algorithm for Experiment 2 to have L1-norm regularization (penalty = 1).

## Experiment 2: Results

By design, the feedback and control non-feedback conditions differed in the overall amount of context-relevant information on the screen. In the non-feedback runs, participants were presented with strong context cues via fully coherent images during the reinstatement phase. On the other hand, in neurofeedback runs, there is overall weaker context evidence on the screen, but there is a link between what the participant sees and what their internal context is. Given these bottom-up perceptual differences, it is perhaps unsurprising that the conditions differed in the total amount of classifier evidence decoded during the context reinstatement period. During the runs with feedback, when participants viewed composite face/scene mixtures, the average accuracy of the multivariate pattern classifier was 0.58, which was reliably above chance (95% CIs 0.49–0.67; chance = 0.5; *p* = < 0.037), as in Experiment 1. During the control runs without feedback, when participants viewed unmixed images, the average accuracy of the multivariate pattern classifier was 0.90 (95% CIs 0.88–0.93; chance = 0.5; p < 0.00001) and this was reliably greater than the feedback runs (p < 0.00001).

The larger goals of this experiment were to (a) replicate the (valid-condition) effects observed in Experiment 1 and (b) investigate whether the relationship between context reinstatement and memory recall was observed in a condition without closed-loop neurofeedback. Our hypothesis was that the link between context reinstatement and memory recall is fostered by our neurofeedback procedure, and therefore would be larger in the feedback condition than in the non-feedback condition. While the intent of the feedback manipulation was to *boost sensitivity* to the brain-behavior relationship (between neural context reinstatement and recall), not to boost overall recall, we nonetheless looked at the effects of the feedback manipulation on recall performance. We found no reliable difference between the average number of words recalled in control, non-feedback runs and neurofeedback runs (recalls_feedback_ = 8.88, 7.75–10.22; recalls_control_ = 9.10, 7.94–10.30, *p* = 0.44; **Figure 5a-c**). It is notable that recall levels were similar across conditions even though scenes were much less visible in the feedback condition. If bottom-up scene perception alone determined recall, we would expect recall to be much better in the non-feedback condition. The fact that recall performance was relatively matched across conditions fits with the idea that top-down context reinstatement also supports recall performance.

**Figure 5.**
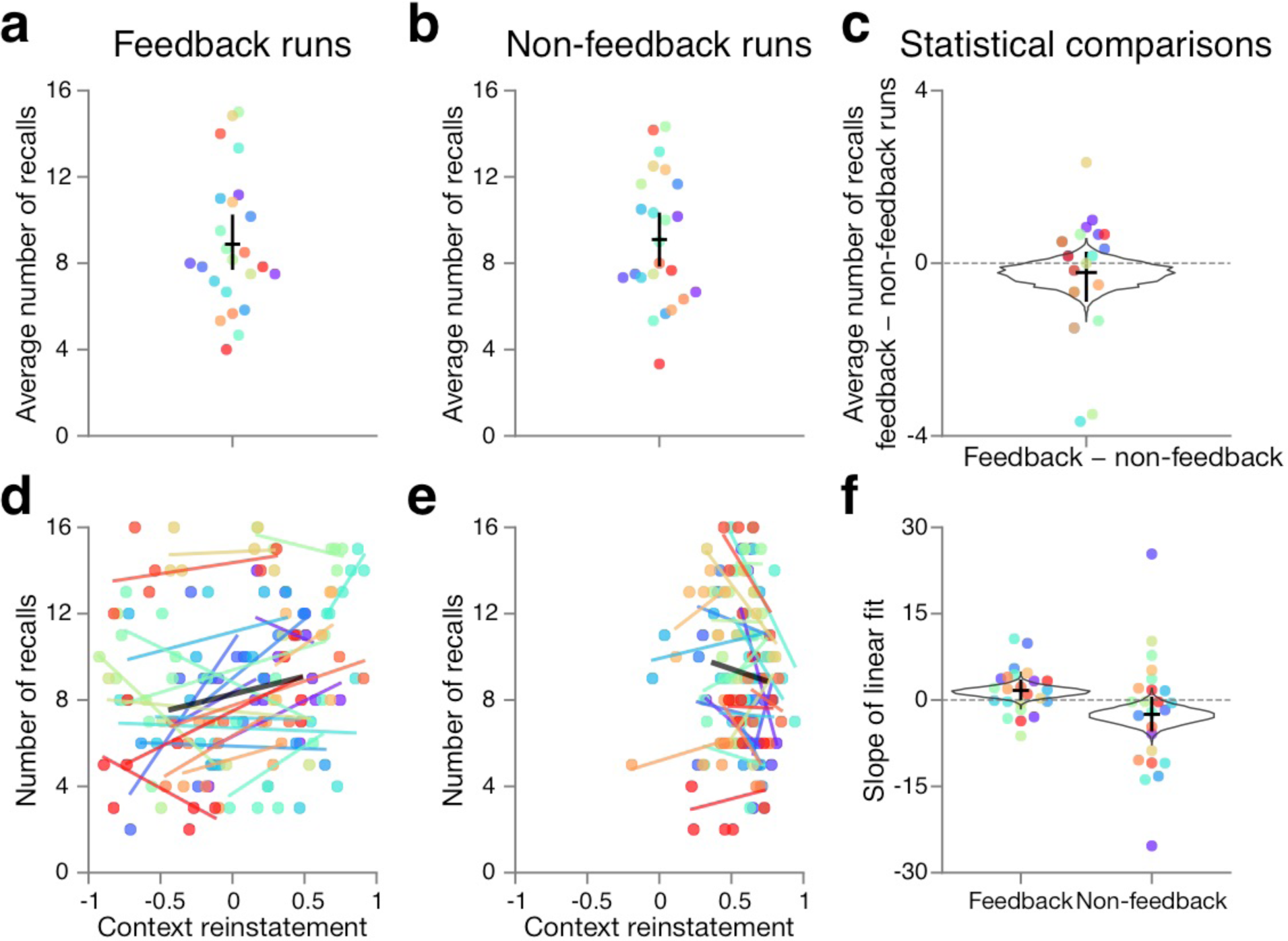
Feedback mediates the link between context reinstatement and memory in Experiment 2. (a) Memory recall performance for validly cued feedback runs. In these runs, feedback was provided during the context reinstatement period. Each dot corresponds to the average number of recalls for a participant (n=24). The horizontal black line indicates the population average, and the vertical black line indicates 95% confidence intervals. (b) Memory recall performance for non-feedback control feedback runs. In these runs, there was no real-time feedback during the context reinstatement period. (c) Memory performance did not reliably differ between these feedback and non-feedback conditions (*p* > 0.1). (d) Evidence for the cued context in the feedback condition was positively related to the number of recalls, replicating the effect in the valid feedback condition of Experiment 1 (*p* < 0.05). The *x,y* position of each dot represents data from a single run. All runs from a participant are depicted in the same color, and the linear fit for that participant is a line in the same color. The black line is the mean fit across the population. (e) Evidence for the cued context in the non-feedback control condition was unrelated to memory recall performance (*p* > 0.1). (f) Statistics were computed using the slopes of the linear fits per condition. Each dot corresponds to the slope of the linear fit in a single condition (feedback or no feedback) for each participant. The relationship between context reinstatement and memory performance was reliably greater in the feedback condition than in the non-feedback condition (*p* < 0.05).

Next, we repeated the analyses developed during the first experiment, to examine whether context reinstatement during feedback runs relates to subsequent memory recall performance. Indeed, we replicated the positive relationship between context reinstatement in validly cued runs and memory performance, this time using an entirely different group of participants (slope*_feedback_* = 1.66, 0.60–3.29 *p* = 0.025; n = 23, 1 outlier excluded, **Figure 5d**).

Then, we investigated whether this same positive relationship was present without neurofeedback. In fact, there was no such relationship between context reinstatement and memory performance in the runs without feedback (slope*_nofeedback_* = −3.47, −7.12 – −0.53; *p* = 0.97; **Figure 5e**); the relationship was actually reliably negative. The difference in slope between the feedback and non-feedback conditions was reliable (*p_diff_* = 0.032). These results fit with our hypothesis that neurofeedback makes it easier to identify a link between context reinstatement and recall performance.

## Discussion

In the studies presented above, we demonstrated that context reinstatement (measured neurally) predicts subsequent free recall success: Reinstating the correct context boosts recall success (Experiment 1, replicated in Experiment 2) and reinstating the incorrect context reduces recall success (Experiment 1). These results extend prior work by Polyn et al. (2005) and others (e.g., Morton et al., 2013), by showing that it is specifically *context* (not item) reinstatement that drives this effect. In this study, contexts and to-be-learned items were different types of stimuli (contexts were face or scene pictures; items were uncategorized words). Furthermore, the neurofeedback was derived from a multivariate pattern classifier that had been trained on an independent localizer period without any words. Taken together, these features of the design make it extremely unlikely that the classifier (applied during the reinstatement period) was picking up directly on recall of words, as opposed to context activation. One intermediate possibility (fully consistent with our theoretical account) is that participants were recalling some words during the reinstatement period, which *caused* context activation (which was then detected by the classifier). However, we think that even this intermediate interpretation is somewhat unlikely: During the reinstatement period, we instructed participants to focus on the images so as to discourage using the reinstatement time to rehearse words. Also, participants were told that they would receive additional monetary reward based on activating the correct context during the reinstatement period, which should have further encouraged them to focus on context reinstatement as opposed to word recall during this period.

Our intent in using neurofeedback was to boost sensitivity to small fluctuations in context reinstatement by amplifying them, thereby making it easier to identify a relationship between context reinstatement (measured neurally) and behavior. That is, we are using neurofeedback to improve measurement sensitivity as opposed to using it as a performance booster. Simulations that we ran (comparing our neurofeedback condition to various controls) support the intuition that neurofeedback can boost experimental sensitivity.

Experiment 2 provides some support for the claim that neurofeedback is especially useful for identifying the relationship between context reinstatement and recall behavior. In this experiment, we compared neurofeedback to a condition where images from original context were 100% visible (we expected that this would be the strongest possible reinstatement cue); we observed a significantly larger relationship between our neural measure of context reinstatement and recall behavior in the neurofeedback condition than in the 100% visible control condition. Previous studies have demonstrated that providing real-time fMRI can reveal insights about cognition (e.g., Cortese et al., 2016; Lorenz et al., 2018). Here, we extend that finding to demonstrate neurofeedback can more tightly link fluctuations of internal mental context with memory retrieval.

Having said this, our conclusions about the specific role of feedback in identifying this relationship are necessarily preliminary. As noted earlier, different control conditions address different issues, and no single control condition can establish that neurofeedback is necessary. The 100%-visible control condition that we used in Experiment 2 may have failed to show an effect because it lacked neurofeedback; alternatively, it may have failed to show an effect for other reasons. For example, there was less variability in classifier evidence in the control condition than the neurofeedback condition — this restricted range effect may have made it harder to link classifier evidence to behavior in this condition. In future work, it will also be highly informative to look at a zero-visibility control condition (i.e., where the screen is blank during the reinstatement period); this control will tell us whether internal mental context reinstatement on its own is sufficient to drive a relationship between brain activity (during the reinstatement period) and recall behavior.

Finally, it is important to emphasize that neurofeedback can be used for multiple purposes: In other, recent neurofeedback studies, feedback has been used to drive learning. For example, this technique has been used for training participants to improve their sustained attention performance (as in deBettencourt et al., 2015), training new associations (as in Amano et al., 2016; see also deBettencourt and Norman, 2016), and reducing established fearful associations (Koizumi et al., 2017). Recently, researchers have used real-time fMRI to link behavior and neural activity, e.g., to dissociate confidence from accuracy (Cortese et al., 2016), to link brain activity with experience in a focused attention task (Garrison et al., 2013), to optimize experimental design (Lorenz et al., 2016), and to characterize a multidimensional task space (Lorenz et al., 2018). Here, we used neurofeedback in a potentially complementary way, to amplify brain activity fluctuations and improve measurement sensitivity for a cognitive process (context reinstatement) that we think is important for memory.

### Conclusions

In this study, we have demonstrated that closed-loop neurofeedback is a useful tool for testing theories of memory retrieval; here, we used it to establish a relationship between context reinstatement (prior to the onset of recall) and memory performance. This technique could be expanded to other experiments in which context has a major role. For example, closed-loop neurofeedback could be used in a task where participants are asked to “flush” context instead of recover context. This process of eliminating context has been demonstrated to have a critical role in intentional forgetting (Manning et al., 2016; Sahakyan and Kelley, 2002). Eventually, it might be possible to further develop this technique to provide training for context reinstatement, and to study and treat psychiatric disorders that involve context-cued recall, such as addiction and post-traumatic stress disorder.

## Acknowledgments

This work was supported by US National Institutes of Health (NIH) grant R01EY021755, US National Science Foundation (NSF) grant BCS1229597, NSF fellowship DGE1148900, the Intel Corporation, and the John Templeton Foundation. The opinions expressed in this publication are those of the authors and do not necessarily reflect the views of these funding agencies.

We would like to thank Paula A. Pacheco for assistance in annotating the recall data, and Anne Mennen and Mariam Aly useful comments on the manuscript. We would also like to thank the PNI computing staff and Intel Labs for technical support related to the real-time analyses.

